# Endozoochory of a dry fruited tree aids quarry passive restoration and seed soaking further increases seedling emergence

**DOI:** 10.1101/561142

**Authors:** Vânia Salgueiro, Carmo Silva, Sofia EufráZio, Pedro A. Salgueiro, Pedro G. Vaz

## Abstract

As practitioners promote passive restoration as a complementary approach to technical reclamation, it is imperative to know its drivers. Although the consequences of endozoochory can be crucial to passive restoration success, few experimental studies assess the use of heavily disturbed sites by seed dispersers such as carnivores and how the seeds they bring in emerge and survive. Using an indoor sowing experiment conducted in a quarry located within a natural park in Portugal, we examined for the first time how carnivore endozoochorous seeds collected in the quarry potentially influence its passive restoration, through effects on plant emergence and survival. Also, we tested whether sowing date and water soaking, relevant factors when sowings are to be carried out, would affect seedling emergence and mortality rates when compared with the effect of endozoochory. Our target species were included in the revegetation plan of the quarry, of which endozoochorous seeds of Carob tree (*Ceratonia siliqua*) were collected in sufficient number for analysis. Irrespective of the carnivore species, endozoochorous carob seeds performed similarly to untreated seeds regarding emergence rates. Endozoochorous carob seedlings showed greater mortality rates but the net result for the plant can still be the colonization of recently vacant habitats by a large proportion of viable seeds. Concerning sowing date, the later carob seeds were sown over the fruit-ripening season, the faster seedlings emerged. Water soaking increased emergence rate by 6.5 times. Broadly, sowings with previous soaking and carnivore-mediated seed dispersal of this dry-fruited tree can jointly enhance quarry restoration.

**Implications for practice:**

- Restorers can undertake pilot sowing experiments prior to large scale revegetation campaigns to identify which species benefit the most from endozoochory.
- Carnivores in the surroundings of a quarry contribute a large proportion of viable seeds, likely assisting quarry passive restoration.
- Carnivores ingesting carob seeds later in the fruiting season may assist quarry passive revegetation more readily as seeds ingested around that time emerge earlier.
- Immersion in tap water is a simple, inexpensive, and highly efficient method to break physical dormancy when carob seed sowings are to be carried out in degraded sites.

## Introduction

A substantial part of current restoration ecology occurs in human-disturbed habitats (Nunes et al. 2016; Prach & Tolvanen 2016). The repair of degraded sites is achieved along a gradient of processes from passive restoration, in which recovery happens through natural succession, to technical reclamation, including plantings and sowings of target species (Prach & Hobbs 2008; Holl & Aide 2011; Cross et al. 2018). The latter dominates in heavily degraded habitats such as quarrying sites but contemporary studies increasingly assert passive restoration as a complementary approach (Prach et al. 2013; Tischew et al. 2014; Sebelikova et al. 2016; Valkó et al. 2017). Assessing potential drivers of passive restoration is thus an urgent task. Several studies investigate the role of substrate quality (Walker & del Moral 2003; van der Putten et al. 2013) and of the surrounding vegetation (e.g., as the source of seeds) on the passive revegetation of heavily degraded sites (Novák & Konvička 2006; Řehounková & Prach 2008; Prach et al. 2015). Type of seed dispersal, seedling emergence, and survival upsurge as major constraints on the revegetation of degraded sites (Jones & del Moral 2009; Zhang & Chu 2015; Horáčková et al. 2016). Few experimental studies, however, document how animal-mediated seed dispersal might influence passive restoration of heavily degraded sites.

The consequences of seed dispersal in animal guts (i.e., endozoochory) to the revegetation of degraded sites are arguably crucial to predict passive restoration success (Cavallero et al. 2013; Horáčková et al. 2016). In topics other than the restoration of heavily degraded sites, research has long been conducted on endozoochorous seed dispersal, especially of fleshy-fruited plants (Herrera 1989; Levey & Benkman 1999; Fedriani & Delibes 2009; Suárez-Esteban et al. 2013; Wotton & McAlpine 2015; García-Cervigón et al. 2018), whereas dispersal responses of dry-fruited plants to endozoochory remain far less studied (but see, e.g., Ramos et al. 2006; Zhou et al. 2013). Because pulp texture affects seed retention time in the gut (Levey 1986), seeds from fleshy- and dry-fruited species may differ in their response to endozoochory and, ultimately, seedling emergence and survival (Traveset & Verdú 2002). Either way, some animal groups provide chief services to plants as endozoochorous seed dispersers (Herrera 2003; Pizo 2007). Carnivores (Order Carnivora), given their large home-ranges and gut-retention times, are suited as dispersal vectors while moving across their territories (Jordano et al. 2007; Koike et al. 2008; Enders & Wall 2012; Herrera et al. 2015; Farris et al. 2017). This role may be species-specific because some carnivores destroy certain seeds while consuming the fruit. Efficient seed dispersers must transport and deposit the seed intact without affecting the viability of the embryo (Rosalino et al. 2010), therefore increasing the likelihood of plant dispersal into degraded sites and boosting passive restoration success.

Determining the net effect of endozoochory on seedling performance (emergence and survival) is challenging, namely in post-quarrying sites characterized by heterogeneous conditions (Clemente et al. 2004; Řehounková & Prach 2006; 2008; Šalek 2013). Seedling performance under field conditions is influenced by a plethora of factors, including substrate quality (e.g., soil or the animal scat itself), deposition site (Milotic & Hoffmann 2016), germination temperature, seed burial depth (Maraghni et al. 2010), germination date (Traba et al. 2006), and microsite moisture. Seed sowing experiments are adequate to avoid confounding effects (Biere 1991; Olmez et al. 2007; Oliveira et al. 2012), especially indoor experiments conducted *in situ*, which reproduce the local photoperiod and light spectrum. In active quarries, indoor sowing experiments may also be preferable to field experiments for safety reasons. In a few related experiments, the effect of endozoochory on seedling performance has been tested (Ortiz et al. 1995; Farris et al. 2017), either using animals kept in captivity (Varela & Bucher 2006) or simulating the gut environment (Milotic & Hoffmann 2016). Endozoochory is seldom examined together with seed water soaking (Tsakaldimi & Ganatsas 2001) and sowing date (Wang & Lechowicz 1998; von Gehren et al. 2016), which are both relevant factors when sowings are to be carried out in degraded sites. Integrated approaches combining the field collection of reliable data on endozoochorous seeds and controlled sowing experiments are needed (Traveset et al. 2007) to better understand the impact of endozoochory on passive restoration.

Herein, we combined intensive field surveys of endozoochorous seeds and an indoor-sowing experiment to address the potential role of carnivore-mediated seed dispersal on quarry passive restoration. In an active quarry within a natural park in Portugal, we set up a sowing experiment with the main objective of comparing the performance (emergence, survival) of seeds dispersed by carnivores into the quarry, seeds previously soaked in water, and untreated seeds. We picked endozoochorous seeds from carnivore scats collected along the quarry paths. Our target-seeds were from the plant species being used by restorers for the artificial revegetation of the quarry. We wonder whether endozoochorous and untreated target-seeds would differ in performance in an *in situ* greenhouse to infer the potential contribution of dispersed seeds to quarry revegetation. In our analyses, the first hypothesis was that carnivore endozoochory would not affect seedling emergence (Ortiz et al. 1995), though passage through the gut could lead to seed mass and water content losses likely decreasing seedling survival later on (Traveset et al. 2008). We expected that endozoochory would still benefit passive revegetation as a proportion of endozoochorous seedlings would potentially colonize new habitats in the quarry. Second, we tested whether water soaking, widely used to accelerate germination in hard-coated seeds (such as Carob tree seeds in our experiment) would cause performance to increase relative to endozoochorous and untreated seeds (Tsakaldimi & Ganatsas 2001; Gunes et al. 2013). Lastly, we tested whether performance would also vary with sowing date, as gradual changes in light and temperature would likely influence the emergence timing throughout the experiment (Morris et al. 2000; von Gehren et al. 2016). By controlling for most factors affecting seed germination in the field, we expect to gain a better understanding of the real implications of endozoochory on quarry restoration.

## Methods

### Study area and seed collection

We conducted this study within a 425-ha property (SECIL-Outão, Setúbal, Portugal), including 99 ha for active limestone and marl quarrying (Fig. 1). SECIL-Outão is located within the Arrábida Natural Park on the eastern slopes of the Arrábida Chain, a mountain area 40 km south of Lisbon, reaching the maximum altitude of 501 m asl. The area has steep slopes and is established on Jurassic limestone formations with Mediterranean red soils (Pedro 1991). The Arrábida Chain has a well-preserved Mediterranean flora with 128 species in vegetation forms varying from shrubland to woodland (Catarino et al. 1982). The climate is Mediterranean sub-humid with marked influence of the Atlantic Ocean. Annual precipitation is 745 mm, falling mainly in fall and winter, while mean annual temperature is 16.8°C, ranging from 2.7°C in December– February to 38.1°C in June–September.

**Figure 1.**
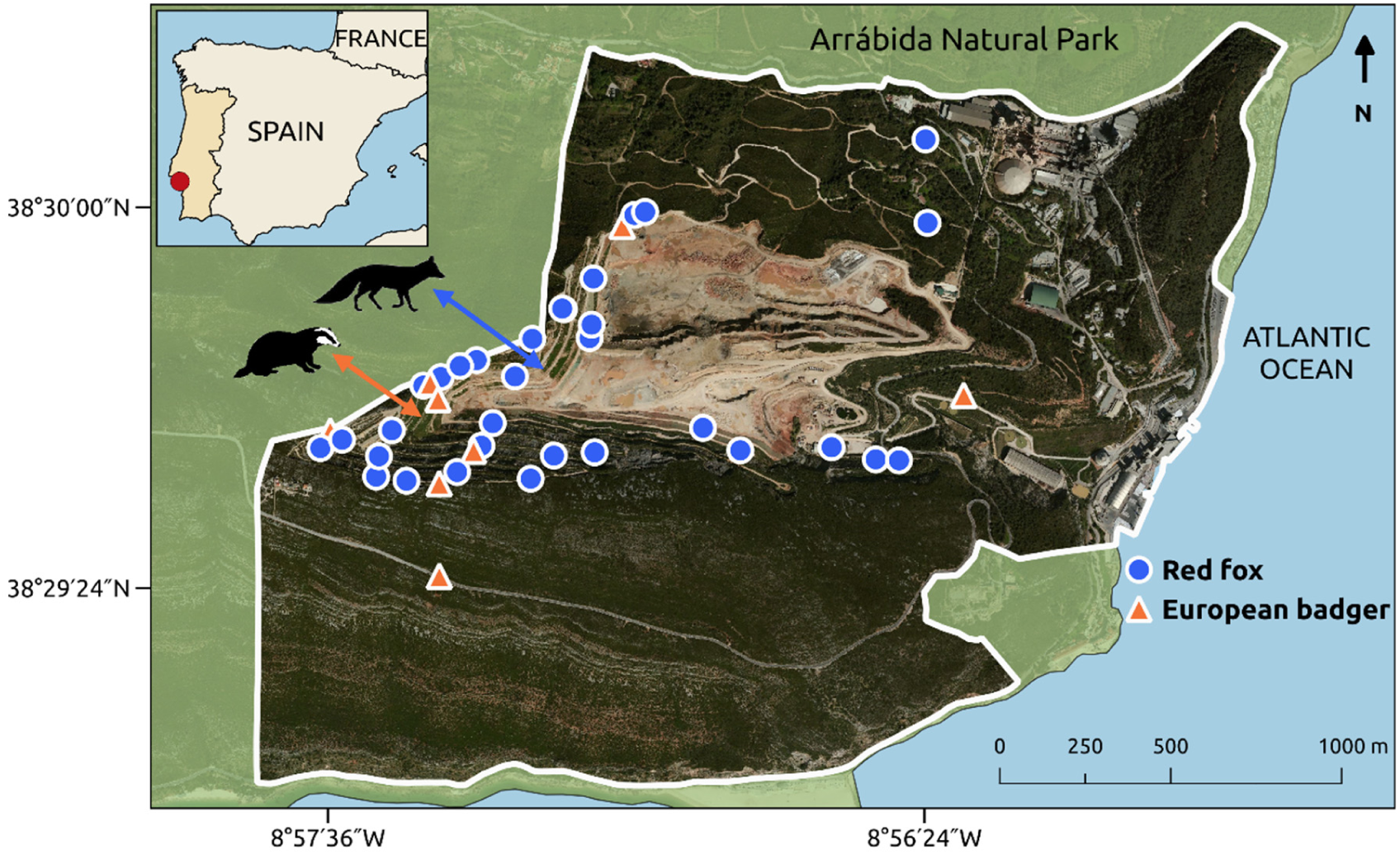
SECIL-Outão limestone quarry area in south-west Portugal and locations of the carnivore scats from which seeds were picked to assess the effect of endozoochory by medium-sized carnivores. Quarry landform (slopes = 35–38°) is noticeable, which include stones, rocky outcrops and crevices.

To accelerate revegetation of vacant terraces after quarrying extraction, SECIL-Outão is restoring those areas with several plant species to foster convergence with the surrounding landscape. This artificial revegetation included indigenous, evergreen species native to the Mediterranean Basin (Clemente et al. 2004). Among these, we would expect Carob tree, Strawberry tree (*Arbutus unedo*), Common myrtle (*Myrtus communis*), and Olive tree (*Olea europaea* var. *sylvestris*) fruits to be common in carnivores’ diet. Conversely, we would expect the remaining species used in the artificial revegetation to either be much less ingested (Herrera 1989; Rosalino et al. 2005; Rosalino & Santos-Reis 2009), or infrequent in carnivore scats, as with Phoenician junipers, *Juniperus phoenicea* (Herrera 1989; but see Farris et al. 2017) and Mastic tree, *Pistacia lentiscus* (Suárez-Esteban et al. 2013). Thus, Carob tree, Strawberry tree, Common myrtle, and Olive tree were the target species for our sowing experiment. The four species were present in the surrounding vegetation in the natural park (Meira-Neto et al. 2011; Espírito-Santo et al. 2011).

Other species that were being used in the artificial revegetation of SECIL-Outão but would be less likely ingested by carnivores according to the literature included Rozeira (*Lavandula luisieri*), Rosemary (*Rosmarinus officinalis*), Broadleaved lavender (*Lavandula latifolia*), Stone pine (*Pinus pinea*), Kerm oak (*Quercus coccifera*), Portuguese oak (*Quercus faginea*), Grey-leaved cistus (*Cistus albidus*), Sage-leaved rock-rose (*Cistus salviifolius*), Evergreen honeysuckle (*Lonicera implexa*), Laurustinus (*Viburnum tinus*), and the Narrow-leaved phillyrea (*Phillyrea angustifolia*). Locally, seeds were potentially ingested and dispersed by the medium-sized carnivore species Red fox (*Vulpes vulpes*), European badger (*Meles meles*), Stone marten (*Martes foina*), Egyptian mongoose (*Herpestes ichneumon*), and Common genet (*Genetta genetta*) (Mira et al. 2014). Because ~40 % of SECIL-Outão perimeter was unfenced and the existing wire fence (≤ 1.4 m height) was installed 20 cm above ground and had large mesh size (15 × 15 cm), carnivores easily access the quarry.

To maximize the collection of endozoochorous seeds, we did an intensive search for carnivore scats from October 2015 to January 2016, locally matching the main fruiting season of our target plant species. Because most medium-sized carnivore species frequently deposit their scats along paths, we focused sampling efforts along the entire 21.1-km network of paths within SECIL-Outão. To control for scat age variation, all paths were cleared of mammal scats one week prior to the beginning of scats’ collection. Thereafter, all paths were searched for scats 16 times, nearly weekly from 2 October 2015 to 26 January 2016. For two consecutive days a week, to reduce inter-observer variability, three trained persons only, two at a time searching through different paths, collected and identified the scats’ species. Whenever there were identification doubts between carnivore species, the scat was recorded as non-identified. If we felt the uncertainty could have arisen from confusion with domestic animals (cats, dogs), the scat was discarded. Immediately after identification at the species level on the basis of shape, odor, color, and place of deposition, we geo-referenced the position and collected each scat into it’s own paper bag. At the end of each sampling day, scats were air dried and stored individually in paper bags in the dark at room temperature. Each scat was later washed using sieves under running water (mesh size = 1 mm). Seeds were immediately and carefully removed and dried. Then, each seed (or group of seeds picked from the same scat) of our target plant species was carefully examined and stored in a paper envelope. We selected only unharmed seeds (i.e., not crushed or fractured) for the sowings. During the scat survey period, local mean daily temperature 4 m above ground was 14.7 °C (7.0–20.6 °C) and the local total precipitation was 503.8 mm (Supporting Information, Fig. S1).

To obtain untreated seeds, plus carob seeds to be later water soaked, we collected ripe fruits in September 2015 from 20–30 individuals of each target species well-distributed throughout the area nearby. Then, we extracted the seeds from the carob pods and from the fleshy-fruits of the other target species (pulp was first removed under running water with a strainer). By species, all viable seeds were then pooled, randomly hand-mixed, and kept dry in the dark at room temperature. We excluded seeds with any sign of pathogens or insect damage. To assign a soaking treatment to carob seeds, they were immersed in tap water at room temperature for one day before sowing.

### Sowing experiment

To test whether emergence and survival would vary with the sowing date over the experiment period (matching the main fruit-ripening season), we did the sowings on 23 November, 9 and 21 December, and 27 January. In each date, all the seeds subject to endozoochory until that moment were sown. The sowing essay was held *in situ* at the SECIL-Outão greenhouse. We used rectangular plastic planting trays (7 × 4 cavities) in which we buried 2 cm deep one seed per cavity filled with commercial soil (cavity width × length × depth = 5 × 5 × 10 cm). Each endozoochorous seed was sown in a cavity contiguous to a cavity with a control seed of the same species. In the case of Carob tree, a third seed, previously soaked, was sown in an adjacent tray cavity. Watering was provided immediately to all seeds after sowing and every three days thereafter. All sowing cavities were daily monitored thereafter for seedling emergence, death, days to emergence, and follow-up days (number of days the seedling was seen alive). We recorded both the last emergence and death events on 9 May 2016 and extended the trial until 1 June 2016. Inside the greenhouse during the monitoring time period, mean daily air temperature (measurements at 8 a.m., noon, and 5 p.m.) was 20.0 °C (10.3–32.8 °C) and the humidity was 55.9 % (27.0–97.7 %). During the sowing trial, air temperatures were on average 4.5 °C higher than outside at the quarry.

### Statistical analysis

Since endozoochory was assessed in seeds that were in some cases picked from the same scat, we first quantified the intraclass correlation (ICC) associated with scat as a grouping variable for seedling emergence and survival. To estimate within-scat ICC for the response variables (seedling emergence, death, days to emergence, and follow-up days), we considered data from scats with two or more seeds only. ICC values below 0.40 would indicate poor degree of within-scat seed resemblance (greater independence of the seeds), whereas 0.40– 0.59 would be fair, 0.60–0.74 would be good, and 0.75–1.00 would be an excellent resemblance (Cicchetti 1994). We used the package *psych* in R to estimate ICC (Revelle 2018). Furthermore, because responses of seeds picked from nearby scats could have been more similar than those sampled at locations more distant from each other, we assed spatial autocorrelation in the endozoochory treatment data subset. We tested the significance of spatial autocorrelation using spline correlograms based on Moran’s *I* (Bjørnstad & Falck 2001) with 95% bootstrap confidence intervals, estimated by the *ncf* package for R (Bjørnstad 2018).

To identify whether seed treatments and sowing date were significant predictors of the emergence or mortality rates of our target plant species, we used two separate Cox regression analyses (Therneau & Grambsch 2000). Per seed, the dependent variable in both survival regressions was calculated from two parts, i.e., the number of days to event (seedling emergence or death) and the event status, which records if the event of interest (emergence or death) occurred or not during the experiment. In both cases, responses were modeled as right-censored due to the uncertainty that seedlings could eventually emerge or die after the experiment. The significance of each predictor was evaluated by backward-stepwise elimination from the full model, which included both seed treatment (levels = control, endozoochory, soaking) and sowing date (levels = 23 November, 9 December, 21 December, 27 January). We assessed a final model for each of the two response variables. In the case of seedling emergence, both explanatory variables were deemed significant (chi-square test; *P* < 0.05), whereas treatment was the only significant predictor in the case of seedling survival regression and so we assessed the final model by dropping sowing date. Before this, still as part of the exploratory analyses, we also used two Cox regressions to assess whether seedling emergence and mortality rates differ by the sole effect of carnivore species in the endozoochory treatment data subset. We fitted all Cox regressions using the *survival* package in R (Therneau 2015).

## Results

Along the total 338 km covered during the search for carnivore scats at the SECIL-Outão quarry (21.1 km × 16 weeks), we collected a total of 242 scats, of which 197 were carnivore scats and 45 were of non-identified species. Among the carnivore scats, 46 scats included seeds of our target plant species, while 164 had seeds of other plant species or had no seeds. Because six carnivore scats (out of 46) were decomposing (e.g., had spreading fungi hyphae) or had seeds already in germination, they were discarded from further analyses. Thus, 40 carnivore scats (32 Red fox and 8 European badger scats) were used for our sowing experiment (Fig. 1).

We sowed a total of 307 seeds (Table 1) in the SECIL-Outão greenhouse, 108 of which were picked from the carnivore scats. Most of these 108 seeds subject to endozoochory by carnivores were Carob tree seeds (91 seeds picked from 39 scats), followed by Common myrtle (nine seeds from three scats), and Strawberry tree seeds (eight picked from two scats). No olive seeds were found. The largest seeds (measured with a digital caliper to the nearest 0.01 mm) were those of carob (diameter = 7.58 mm ± 0.09 SE, length = 9.60 mm ± 0.10 SE), followed by myrtle (2.60 mm ± 0.08, 3.42 mm ± 0.14) and strawberry (1.37 mm ± 0.05, 2.73 mm ± 0.07). About 72 % of sown seeds emerged, of which 86 % survived by the end of the study period. Owing to insufficient sample size, we excluded myrtle and strawberry seeds from our statistical analyzes of emergence and mortality rates.

**Table 1.** Percentage of emergence, mean time to emergence (± SE), percentage of deaths, and mean follow-up time (± SE) of seedlings that underwent treatments at the SECIL-Outão greenhouse (SW Portugal). Endozoochory = seeds picked from carnivore scats collected across the SECIL-Outão study area. Soaking (Carob tree only) = seeds collected from local plants within one week prior, randomly mixed, and immersed in tap water at room temperature for one day before sowing. No treatment = untreated seeds collected from local plants.

| Species | Treatment | *n* | Emergences | Days to emergence | Deaths | Follow-up days |
| --- | --- | --- | --- | --- | --- | --- |
| Carob tree | No treatment | 91 | 80 % (73) | 49.7 ± 4.1 | 11 % (8) | 103.1 ± 4.9 |
| | Endozoochory | 91 | 54 % (49) | 50.8 ± 5.7 | 20 % (10) | 87.5 ± 7.4 |
| | Soaking | 91 | 91 % (83) | 21.1 ± 1.4 | 8 % (7) | 135.3 ± 4.6 |
| Common myrtle | No treatment | 9 | 44 % (4) | 35.5 ± 1.0 | 0 % | 82.0 ± 22.5 |
| | Endozoochory | 9 | 22 % (2) | 35.0 ± 2.0 | 0 % | 114.5 ± 22.5 |
| Strawberry tree | No treatment | 8 | 63 % (5) | 52.8 ± 6.4 | 60 % (3) | 85.6 ± 24.0 |
| | Endozoochory | 8 | 63 % (5) | 50.8 ± 4.0 | 60 % (3) | 100.0 ± 12.1 |

The ICC associated with scat as a grouping variable was poor for seedling emergence (ICC1 = 0.08) and seedling death (ICC1 = 0.03). Likewise, ICC associated with scat was poor for the number of days from sowing to emergence (ICC1 = 0.36) and for seedling follow-up days (ICC1 = 0.11). Seedling emergence and survival both did not significantly differ (Type-III chi-square test, *P* > 0.05) by carnivore species (European badger or Red fox). The analysis of spline correlograms showed there was no significant spatial autocorrelation in seedling emergence, days to emergence, seedling death, and follow-up days up to a distance of 2 km among scats (Supporting information, Fig. S2). Considering these analyses, we did not distinguish between the two carnivore species and assumed the independence of all seeds subject to endozoochory in subsequent analyzes.

### Effects on seedling emergence

Cox regression analysis indicated seedling emergence of Carob tree seeds was significantly affected (*χ*^*2*^ = 55.23, df = 2, *P* < 0.0001; Wald chi-square test) by seed treatment (Supporting Information, Table S1). Water soaking increased emergence rate by 6.5 times (Fig. 2A). While seedling emergence by previously soaked seeds occurred mostly between the first and fourth week after sowing, control seeds and those that underwent carnivore endozoochory emerged until the 21^st^ week after sowing (Fig. 2B). Seedling emergence varied with sowing date as well (*χ*^*2*^ = 46.09, df = 3, *P* < 0.001). Seeds sown on 9 December, 21 December, and on 27 January had emergence rates 1.6, 2.4, and 3.5 times greater than that of seeds sown on 23 November (Fig. 2C). In general, seeds sown later took less time to emerge (Fig. 2D).

**Figure 2.**
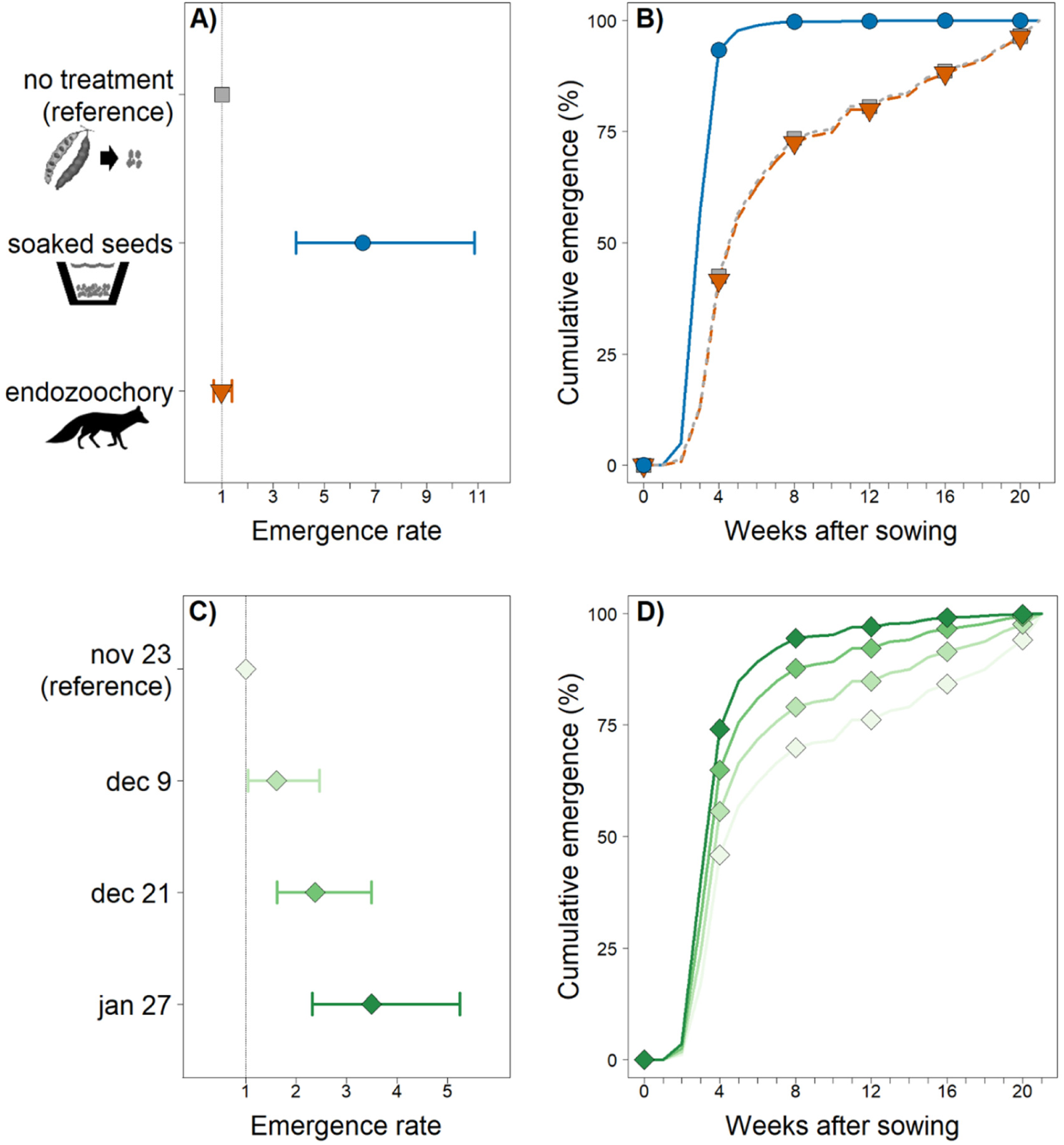
Effect size with 95 % CI (panels A, C) and adjusted cumulative emergence curves (B, D) for the optimal Cox regression model predicting emergence rate of Carob tree seedlings. Endozoochory = seeds picked from carnivore scats. Soaked seeds = immersed in tap water before sowing. No treatment = untreated before sowing (control). Nov 23, Dec 9, Dec 21, and Jan 27 = sowing dates. In panels A and C, vertical dashed lines indicate a rate of 1 (emergence rate similar to the reference).

### Effects on seedling survival

Cox regression revealed that seed treatment significantly affected (*χ*^*2*^ = 7.0, df = 2, *P* = 0.030) seedling mortality rates (Table S2). Seedlings whose seeds underwent endozoochory had a mortality rate 2.1 times greater than that of control seedlings (Fig. 3A). For example, by the fourth week after emergence, the survival proportion of seedlings whose seeds underwent endozoochory was 0.83, while seedlings whose seeds were subject to water soaking or untreated had 0.96 and 0.93 survival proportion, respectively (Fig. 3B). In all three treatments, the survival proportion dropped to its minimum at 16 weeks after emergence, reaching 0.77, 0.91, and 0.88 for endozoochory, soaking, and control treatments.

**Figure 3.**
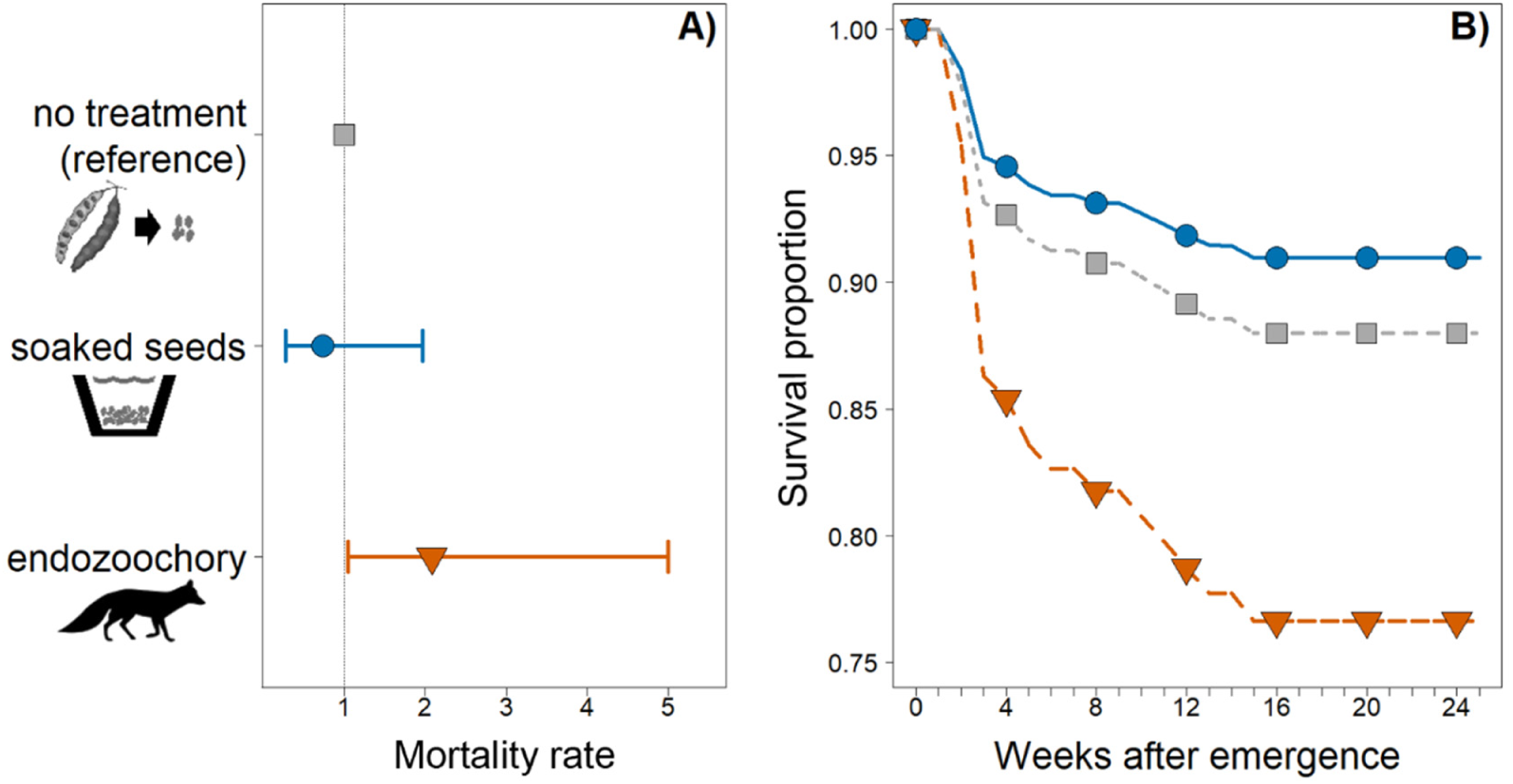
Effect size (A) with 95 % CI and adjusted survival curves (B) for the optimal Cox regression model predicting survival of Carob tree seedlings. Endozoochory = seeds picked from carnivore scats. Soaked seeds = immersed in tap water before sowing. No treatment = untreated before sowing (control). In panel A, the vertical dashed line indicate a rate of 1 (mortality rate similar to control seeds).

## Discussion

Few studies assess the use of heavily disturbed sites such as quarries by terrestrial mammals and their contribution to post-quarrying revegetation via endozoochorous seed dispersal. Using an indoor sowing experiment conducted *in situ*, we examined for the first time whether and how carnivore endozoochorous seeds collected in a quarry potentially influence its passive restoration, through effects on plant emergence and early survival. Most interesting in our analysis were the results for Carob tree, an evergreen species with conservation and socio-economic value (Zohary 2002; Ramon-Laca & Mabberley 2004; Baumel et al. 2018) being used for the artificial revegetation of the quarry. With likely positive consequences for passive restoration, we found that, irrespective of the carnivore species (Red fox or European badger), endozoochorous carob seeds collected in the quarry performed similarly to untreated local seeds regarding emergence rates. Also, although endozoochorous carob seedlings showed greater mortality rates, the net result for the plant can still be the colonization of recently vacant habitats by a large proportion of viable seeds. Namely because plantings and sowings can be expensive (Schirmer & Field 2000; Prach & Pyšek 2001; Cross et al. 2018), we urge restorers elsewhere to undertake similar pilot sowing experiments prior to large-scale revegetation campaigns and thus identifying which species can benefit the most from endozoochory, provided that adequate microsites are present to maximize passive revegetation success.

### Endozoochory of a dry-fruited tree into a quarry

Because dispersal responses of dry-fruited species to endozoochory are less studied than those of fleshy-fruited species, our Carob tree results are further noteworthy. Carob pods are 10–30 cm long dry fruits (i.e., having a dry, hard pericarp) that do not open to release seeds when mature. Favoring the endozoochory mechanism, carnivores jointly ingest seeds, rich in protein, and the pod, very rich in sugars (El-Shatnawi & Ereifej 2001). Carob pods have been important in the diet of farm animals and have been eaten by humans throughout history in the Mediterranean (Zohary 2002; Tous et al. 2013). Larger fruit size and high edibility (Papaefstathiou et al. 2018) may have been reasons carob seeds were far more common in carnivore scats at the SECIL-Outão quarry than seeds from Strawberry tree and Common myrtle as well as olives that were not found. Indeed, despite our intensive sampling effort, we could not gather an acceptable number of carnivore-ingested seeds of these fleshy-fruited species. Still, seeds of Strawberry tree, Common myrtle, and Olive tree were frequent in scats of these carnivore species in ecological studies elsewhere (Kruuk & Kock 1981; Herrera 1989; Aronne & Russo 1997; Santos et al. 2007; Rosalino & Santos-Reis 2009). It remains possible that carnivore endozoochory of these three species benefit quarry passive restoration in other Mediterranean heavily degraded sites. Moreover, although beyond the scope of this study, the large percentage of scats with other than our target seed species indicates that carnivore endozoochory was providing an opportunity for dispersal and possible colonization of the quarry by other plant species as well (Rost et al. 2012; Tischew et al. 2014; López-Bao et al. 2015).

### Performance of endozoochorous seeds

Focusing on the effects on Carob tree, our hypothesis that endozoochory by carnivores of the quarry surroundings would not affect seedling emergence significantly relative to untreated seeds was supported. For endozoochorous seeds, in line with Ortiz et al.’s (1995) results for fox and cow in southern Spain, our data showed that endozoochory decreased percent seed emergence but the rate of decrease was not deemed significant. Our results are also more compatible with no important effect of carnivore species on the performance of endozoochorous seeds. However, despite our high experience in identifying carnivore signs in the field (e.g., Craveiro et al. 2019), it is important to note that there is an uncertainty associated with the scat identification. Several authors suggest the use of molecular approaches to correct known bias associated with researcher experience and scat morphological variations (Monterroso et al. 2013).

Our 80 % emergence for carob untreated seeds is higher than that in most previous studies under controlled situations comparing carob seeds subject to different treatments with untreated seeds. Indeed, related works used Petri dishes for germination trials in laboratory, reaching ≤ 25 % emergence (Ortiz et al. 1995; Tsakaldimi & Ganatsas 2001; Pérez-García 2009; Gunes et al. 2013). Further, because plant provenance has a strong effect on revegetation success due to processes such as local adaptation (see Hufford & Mazer 2003 review), our use of untreated carob seeds collected nearby likely had a positive effect on emergence rate (Martins-Loução et al. 1996). Testing the importance of seed provenance in seedling performance will require further experimental studies (Fedriani et al. 2019). On the other hand, Ortiz et al. (1995) stress that Red fox carry out endozoochorous dispersal of carob seeds without accelerating germination but also not damaging the embryo. Owing to a physical dormancy enhanced by a hard coat, carob seeds can persist viable for several years in the soil (Baskin & Baskin 1989; Ortiz et al. 1995). Because dormancy varies among seeds, dormancy-breaking (e.g., via seasonal temperatures) might be delayed (Pérez-García 2009, Jaganathan et al. 2016). Thus, despite 54 % of our endozoochorous carob seeds having emerged, this percentage could still have increased after the experiment terminus, therefore further potentially benefiting the passive colonization of the quarry.

As hypothesized, the early survival of carob seedlings was diminished by prior endozoochorous seed dispersal. A possible explanation for this is that mechanical and chemical abrasion of the seed coat when passing through the carnivore gut may have had detrimental effects on later seedling condition and survival. Changes in seed mass, water content, permeability, and resistance were previously documented in other common Mediterranean plants (Traveset et al. 2008). Yet, it is worth noting that 77 % endozoochorous seedlings survived our sowing experiment, which is rather promising for the potential colonization of new habitats in vacant terraces after quarrying extraction. However, because sowing experiment conditions (laboratory, greenhouse, field) are of great importance in determining germination (Robertson et al. 2006; Traveset et al. 2007; Fedriani & Delibes 2009) and, ultimately, seedling fate, extrapolating our results to field situations must be considered cautiously. Although the conclusions of our experimental study are certainly valuable to guide revegetation actions taking advantage of the endozoochory mechanism, our results call for further evaluations under heavily degraded field conditions. Namely, whereas we have provided appropriate substrate for our sowings, suitable soil for species such as carob trees may not be readily available in recent vacant terraces after quarrying extraction. Indeed, when more immediate results are required (e.g., to reduce erosion effects), procedures such as hydroseeding and sowings with previous soaking (see below) may be effective (Oliveira et al. 2012). Also, since our experiment ended in June, not covering the driest period of the year, more seedlings could have died due to desiccation over that period, likely accounting for a reduction across treatments in the survival recorded in our results. Because survival to the first summer is an ecological filter regulating plant populations in the Mediterranean (Pugnaire & Valladares 2007), further research on later carob life stages spanning that critical filter is also needed.

The effect of endozoochory on passive restoration of degraded areas depends also on how the scats are delivered in the species’ territory. Many seed-dispersing mammals such as the Red fox, with more scats collected in our study, seem to positively select road verges for defecation (Rost et al. 2012) in a relatively scattered manner (Lloyd 1980), potentially promoting roadside passive restoration (Suárez-Esteban et al. 2013). Although sampling in our work has focused on the quarry paths because they are frequently used by most medium-sized carnivores to defecate, the efficacy of endozoochory will depend on dispersal into degraded areas away from paths too. In the case of European badger, scats may be delivered clustered in shared small dugs (latrines) often buried shallowly, implying the natural sowing of endozoochorous seeds, which may be located far from paths in the degraded area (e.g., along territory boundaries; Kruuk 1989; Revilla & Palomares 2002). Some authors suggest that badgers are among the most efficient seed dispersers (Fedriani & Delibes 2009).

### Sowing date and seed water soaking

The later carob seeds were sown, the faster seedlings emerged across treatments. Temperature and light conditions from November to first fortnight of February were likely important to increasingly alleviate dormancy and trigger emergence, as most other abiotic (e.g., soil moisture) and biotic (e.g., plant competition, insect herbivory) drivers of emergence were constant during the experiment (Forcella et al. 2000, Vaz et al. 2019). Although not a primary objective of the study, our ability to draw inferences regarding optimal emergence timing would have been stronger had it been possible to sow seeds after January. Yet, because carob fruit-ripening season had begun in September, the emergence timing recorded in our study challenges the idea that large-seeded plants (as is the case of Carob tree; Ortiz et al. 1995, Tavsanoglu & Pausas 2018) may tend to emerge early and grow for a longer period (Verdú & Traveset 2005). The extended growth time span can be enhanced by reserves stored in the seed and ultimately benefit seedling survival (Westoby et al. 2002; but see Moles & Westoby 2004). Further, early emergence is in general presumed to be particularly advantageous for perennial Mediterranean species, growing sufficiently during spring to survive summer (Verdú & Traveset 2005). However, early-emerged seedlings may also have higher mortality risk due to seasonal hazards (pathogens, predation, desiccation; Verdú & Traveset 2005). Our results indicate that carob seed sowings by late January, five months before the Euro-Mediterranean summer, are seemingly more advantageous from the restorer perspective than earlier sowings, favoring readily seedling emergences. While for seeds sown earlier the emergence time may not be as critical yet, for seeds sown later a faster emergence allows the seedlings to grow over a longer period before summer.

Our study clearly demonstrates the strong effect of water soaking leading to much faster carob seedling emergence. This information is certainly important when sowings are to be conducted, e.g., as part of a technical reclamation approach. Previous studies evaluating the effect of water soaking on carob emergence used Petri dishes for germination trials in laboratory and seeds previously immersed for a few minutes in hot water (Tsakaldimi & Ganatsas 2001; Pérez-García 2009; Gunes et al. 2013). They followed the same trend, in which water soaking appeared to be a determinant for faster emergence, even though using different methods. Pérez-García (2009) further tested the effect of soaking in distilled water at room temperature on germination and did not find a significant effect. Our water soaked seeds, immersed in tap water at room temperature for one day before sowing, had the highest final percentage of emergence compared to previous studies (91 %). We propose local restorers efficiently undertake breaking physical dormancy in this species by immersion in tap water as it seems to be a simple, inexpensive, and highly efficient method.

Empirical evidence in our study revealed that carnivores in the surroundings of a quarry can contribute a large proportion of viable seeds, likely assisting quarry passive restoration. In the case of Carob tree, endozoochory appears to not affect seed emergence rate and, although increasing seedling hazard rate, a large proportion of seedlings can still survive. The carnivore contribution to the quarry passive restoration can vary throughout the fruit-ripening season. From our case study, carnivores ingesting carob seeds later in the fruiting season may assist quarry passive revegetation more readily as seeds ingested around that time emerge earlier. Further, whenever seed sowing is to be carried out following quarrying extraction, we show that, as expected, water soaking implies faster emergences for carob seeds, although that may not affect seedling survival. Broadly, sowings with previous soaking, and carnivore-mediated seed dispersal of this dry-fruited tree can jointly enhance quarry restoration.

## Acknowledgements

Fieldwork and logistics supported by SECIL-Companhia Geral de Cal e Cimento, SA (project “Estudo e Valorização da Biodiversidade – Componente da Fauna – na Propriedade SECIL-Outão”). Fundação para a Ciência e a Tecnologia funded PGV (SFRH/BPD/105632/2015), and the indirect costs (overheads) of CEABN-InBIO (UID/BIA/50027/2013 and UID/BIA/50027/2019). We are especially indebted to Eulália Freitas and Alexandra Silva for their invaluable assistance in the experiment, and to Ana Machado and João Craveiro for assistance in the processing of samples in the laboratory.

## Supporting information

**Figure S1.**
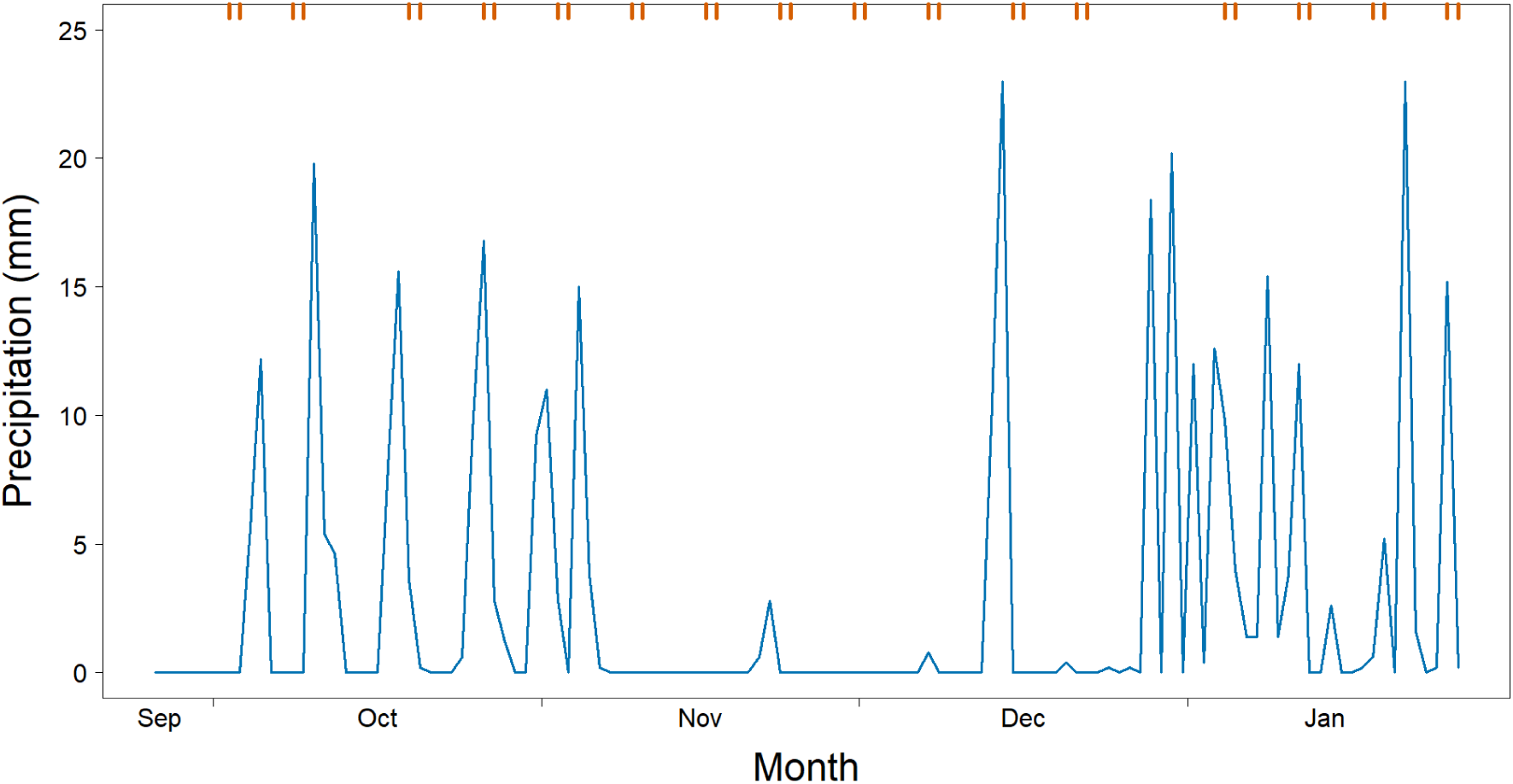
Daily precipitation during the period of collection of carnivore scats (2015–2016) at the SECIL-Outão limestone quarry, south-west Portugal. Upper axis tick marks = days in which scats were collected along the network of paths surrounding the quarry area.

**Figure S2.**
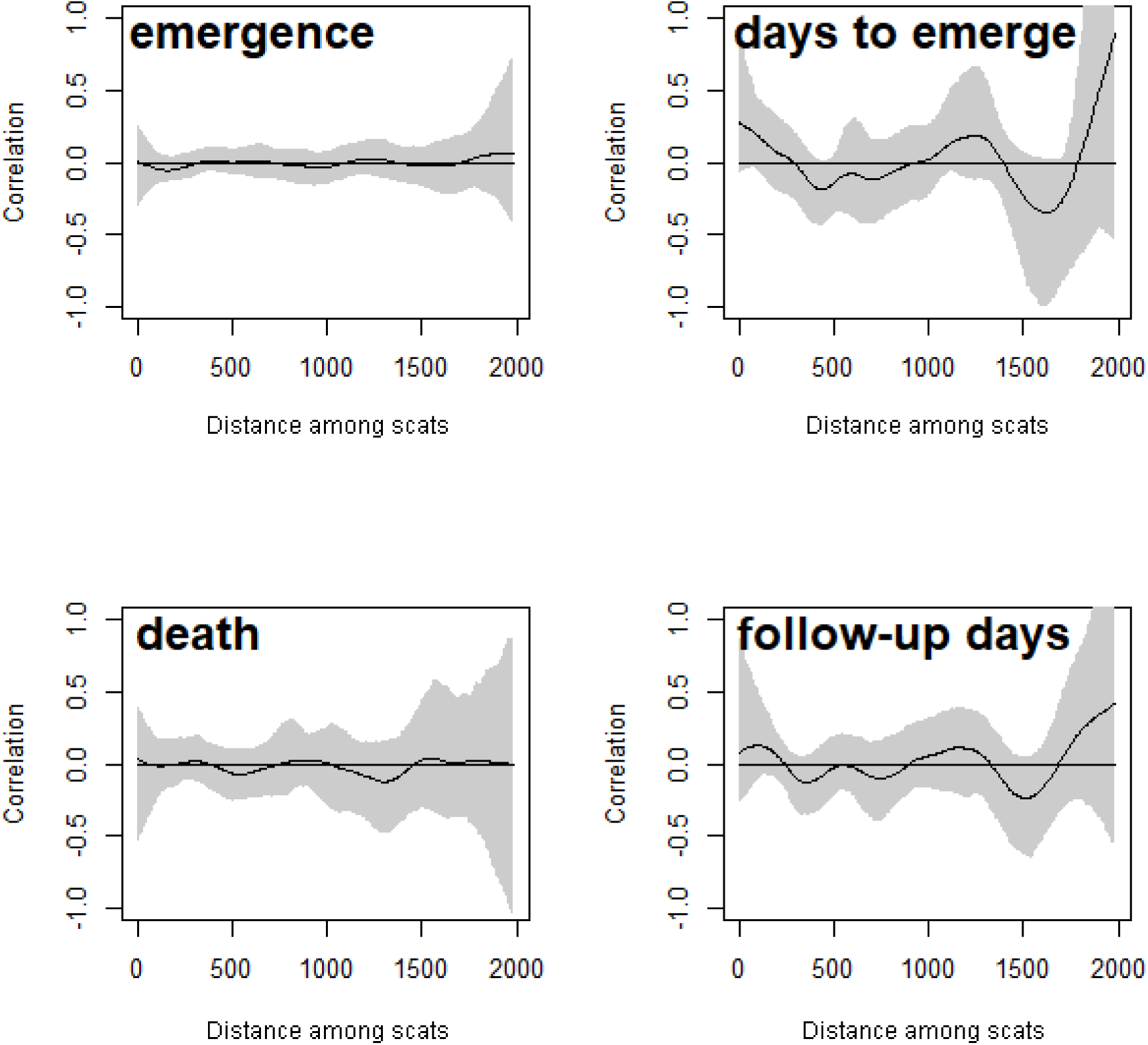
Spline correlograms (Moran’s *I*), with 95 % bootstrap confidence intervals, of Carob tree seeds picked from carnivore scats at the SECIL limestone quarry, south-west Portugal. Spatial correlations of seedling emergence events, days to emergence, seedling death events, and follow-up days assigned to the location of the respective scats are assessed.

**Table S1.** Survival model by Cox regression for emergence rate of Carob tree seeds. HR = hazard (emergence) rate relative to the variable reference level (control seeds for seed treatment and November 23 for sowing date).

| Wald $\chi^2 = 148.6$, $P < 0.0001$; concordance = 0.77 | | | | | |
| --- | --- | --- | --- | --- | --- |
| Variable | β-coef. | SE | $z$ | $P$ | HR (95%CI) |
| seed treatment (soaked seeds) | 1.872 | 0.192 | 7.13 | <0.0001 | 6.50 (3.89–10.88) |
| (endozoochory) | -0.034 | 0.188 | -0.19 | 0.850 | 0.97 (0.68–1.38) |
| sowing date (December 9) | 0.474 | 0.207 | 2.18 | 0.029 | 1.61 (1.05–2.46) |
| (December 21) | 0.869 | 0.193 | 4.49 | <0.0001 | 2.38 (1.63–3.49) |
| (January 27) | 1.249 | 0.214 | 6.00 | <0.0001 | 3.49 (2.32–5.24) |

**Table S2.** Survival model by Cox regression for the mortality rate of Carob tree seedlings. HR = hazard rate relative to control seeds (untreated).

| Variable | Wald $\chi^2 = 6.8$, $P = 0.03$; concordance = 0.61 | | | | |
| --- | --- | --- | --- | --- | --- |
| | β-coef. | SE | $z$ | $P$ | HR (95%CI) |
| seed treatment (soaked seeds) | -0.306 | 0.518 | -0.61 | 0.542 | 0.74 (0.28–1.97) |
| (endozoochory) | 0.731 | 0.475 | 2.05 | 0.041 | 2.08 (1.05–5.00) |

